# CIITA induces expression of MHC-I and MHC-II in transmissible cancers

**DOI:** 10.1101/2021.07.14.452299

**Authors:** Chrissie E. B. Ong, Yuanyuan Cheng, Hannah V. Siddle, A. Bruce Lyons, Gregory M. Woods, Andrew S. Flies

## Abstract

MHC-I and MHC-II molecules are critical components of antigen presentation and T cell immunity to pathogens and cancer. The two monoclonal transmissible devil facial tumours (DFT1, DFT2) exploit MHC-I pathways to overcome immunological anti-tumour and allogeneic barriers. This exploitation underpins the ongoing transmission of DFT cells across the wild Tasmanian devil population. We have previously shown that constitutive expression of NLRC5 can induce stable upregulation of MHC-I on DFT1 and DFT2 cells, but unlike IFNG-treated cells, NLRC5 does not upregulate PDL1. MHC-II expression is crucial for CD4^+^ T cell activation and is primarily confined to haematopoietic antigen presenting cells. Transcriptomic analysis of DFT1 and DFT2 cell lines showed that several genes of the MHC-I and MHC-II pathways were upregulated in response to constitutive overexpression of the class II transactivator (CIITA) gene. This was further supported by upregulation of MHC-I protein on DFT1 and DFT2 cells, but interestingly MHC-II protein was upregulated only on DFT1 cells. The functional significance of the MHC upregulation on DFT cells was shown using serum from devils with natural or immunotherapy-induced DFT1 regressions; binding of serum IgG was stronger in CIITA-transfected cells than wild type cells, but was less than binding to NLRC5 transfected cells. This new insight into regulation of MHC-I and MHC-II in cells that naturally overcome allogeneic barriers can inform vaccine, immunotherapy, and tissue transplant strategies for human and veterinary medicine.

## 1. Introduction

The Tasmanian devil is the largest extant carnivorous marsupial and is endemic to the island state of Tasmania. Following the emergence of devil facial tumour disease (DFTD) in 1996, the population of devils has declined precipitously, threatening the persistence of devils in the wild^1^. DFTD is caused by two independent transmissible cancers of Schwann cell origin, referred herein as DFT1 and DFT2^2,3^. DFT1 was discovered northeast of Tasmania in 1996 while the second tumour, DFT2, was found in 2014 in the D’Entrecasteaux channel, southeast Tasmania. Both tumour types are clonal cell lines that harbour distinct genetic profiles differing from individual host devils^2,3^. DFT cells are transmitted as a malignant allograft amongst devils through social interactions.

Genetic differences between host and tumour, particularly at the major histocompatibility complex (MHC) loci^4^, should induce immune-mediated allograft rejection. However, the 25 years of ongoing DFT1 transmission suggests that DFT1 cells have evolved to evade immune defences. The lack of anti-DFT immune responses has predominantly focused on the loss of MHC-I from the surface of DFT1 cells. This occurs via epigenetic downregulation of several components of the MHC-I antigen processing pathway^5^ and a hemizygous deletion of beta-2 microglobulin (*B2M*), which is necessary for stabilising MHC-I complexes on the cell surface^6^. Natural and immunotherapy-induced tumour regressions have been observed in devils, along with antibody responses to DFT1 cells, albeit primarily in the context of MHC-I^7–9^. Conversely, the emerging DFT2 tumours do express MHC-I^10^, suggesting that other immune evasion mechanisms are important.

Given the role of MHC-I in antigen display and anti-DFT humoral response, the manipulation of MHC-I expression on DFT cells is an attractive target to improve host responses towards DFT cells and mitigate the effects of disease in the wild devil population. An upregulation of MHC-I on DFT cells should enhance MHC-I-restricted tumour-specific cytotoxic CD8^+^ T cell response. However, this approach alone proved to be insufficient for eliciting protective immunity, as exemplified in immunisation trials of naïve devils against DFT1^9^. Although CD8^+^ T cells are recognised as the major effector cells in tumour elimination, CD4^+^ T cell help is critical in facilitating an effective anti-tumour immune response. CD4^+^ helper T cells play a multifaceted role of orchestrating the adaptive and humoral immune response. From cytokine production to expression of co-stimulatory molecules, CD4^+^ helper T cells initiate, augment, and sustain the effector function of not only CD8^+^ T cells and B cells but also innate cells^11–14^. Moreover, CD4^+^ T cells are capable of initiating allograft rejection independently of CD8^+^ T cells^15,16^.

The activation of CD4^+^ T cells involves recognition of antigens presented on MHC-II complexes. In contrast to MHC-I, constitutive expression of MHC-II is restricted to thymic epithelial cells, activated human T cells, and professional antigen presenting cells (APCs) such as B cells, dendritic cells, and macrophages. However, *de novo* MHC-II expression can be induced in non-haematopoietic cells including tumour cells by the inflammatory cytokine interferon gamma (IFNG)^17^. Both constitutive and IFNG-induced expression of MHC-II genes are mediated by the Class II transactivator (CIITA), making it the master regulator of MHC-II expression^18,19^. Additionally, CIITA is capable of modulating the expression of MHC-I, particularly in cell lines with low to no MHC-I expression^20,21^.

The presence of MHC-II molecules in DFT cells has not been described, although CIITA and some MHC-II transcripts can be upregulated *in vitro* in DFT1 cells with IFNG treatment^5^. We have previously genetically modified DFT1 and DFT2 cells that overexpress the MHC-I transactivator NLRC5 to induce stable expression of MHC-I on the cell surface^8^. The lack of MHC-II expression in DFT cells provided an opportunity to conduct similar investigations into the role of CIITA in MHC-II regulation in marsupials and transmissible cancers. Transcriptomic and protein-based analyses showed that CIITA upregulates the expression of genes associated with MHC-I and MHC-II antigen processing and presentation in DFT cells. The ability to modulate antigen presentation in transmissible cancer cells in the context of MHC uncovers additional targets for anti-tumour immune response and the potential for recruitment of CD4^+^ T cell help.

## 2. Materials and methods

### 2.1 Cells and cell culture conditions

Cell lines that were used in this study include DFT1 cell line C5065 strain 3^22^ (RRID:CVCL_LB79), and DFT2 cell lines: RV (RRID:CVCL_LB80) and JV (RRID:CVCL_A1TN)^3^ (**Table 1**). DFT1 C5065 was provided by A-M Pearse and K. Swift of the Department of Primary Industries, Parks, Water and Environment (DPIPWE) (Hobart, TAS, Australia) and was previously established from DFT1 biopsies obtained under the approval of the Animal Ethics Committee of the Tasmanian Parks and Wildlife Service (permit numbers A0017090 and A0017550)^22^. DFT2 cell lines RV and JV were established from single cell suspensions obtained from tumour biopsies^3^. Cells were cultured at 35 °C with 5% CO_2_ in Gibco™ RPMI 1640 medium with L-glutamine (Thermo Fisher Scientific, Waltham, MA, USA) supplemented with 10% heat-inactivated fetal bovine serum (Bovogen Biologicals, Melbourne, VIC, Australia), 1% (v/v) Gibco™ Antibiotic-Antimycotic (100X) (Thermo Fisher Scientific), 10 mM Gibco™ HEPES (Thermo Fisher Scientific) and 50 μM 2-mercaptoethanol (Sigma-Aldrich, St. Louis, MO, USA) (complete RPMI medium).

**Table 1.** List of all devil facial tumour (DFT) cell lines and treatments

| ID # | Sample name | Parent cell line | Treatment | References | ENA project # |
| --- | --- | --- | --- | --- | --- |
| 1 | DFT1.WT | DFT1 C5065 | Untreated | Patchett et al., (2018) | PRJNA416378 |
| 2 | DFT2.WT<sup>RV</sup> | DFT2 RV | Untreated | Patchett et al., (2020) | PRJEB28680 |
| 3 | DFT2.WT | DFT2 JV | Untreated | Ong et al., (2021) | PRJEB39847 |
| 4 | DFT1.WT + IFNG | DFT1 C5065 | 5 ng/mL IFNG, 24h | Ong et al., (2021) | PRJEB39847 |
| 5 | DFT2.WT<sup>RV</sup> + IFNG | DFT2 RV | 5 ng/mL IFNG, 24h | Ong et al., (2021) | PRJEB39847 |
| 6 | DFT1.BFP | DFT1 C5065 | Transfected with empty vector pSBbi-BH | Ong et al., (2021) | PRJEB39847 |
| 7 | DFT2.BFP | DFT2 JV | Transfected with empty vector pSBbi-BH | Ong et al., (2021) | PRJEB39847 |
| 8 | DFT1.NLRC5 | DFT1 C5065 | Transfected with NLRC5 vector pCO1 | Ong et al., (2021) | PRJEB39847 |
| 9 | DFT2.NLRC5 | DFT2 JV | Transfected with NLRC5 vector pCO1 | Ong et al., (2021) | PRJEB39847 |
| 10 | DFT1.CIITA | DFT1 C5065 | Transfected with CIITA vector pCO2 | This study | PRJEB45867 |
| 11 | DFT2.CIITA | DFT2 JV | Transfected with CIITA vector pCO2 | This study | PRJEB45867 |

### 2.2 Plasmid construction

The coding sequence for full length devil *CIITA* (XM_023497584.2) was isolated from cDNA of devil peripheral blood mononuclear cells (PBMCs) by PCR using Q5^®^ Hotstart High-Fidelity 2X Master Mix (New England Biolabs (NEB), Ipswich, MA, USA) (see **Supplementary Table 1** for list of primers and reaction conditions). Sleeping Beauty (SB) transposon plasmid pSBbi-BH^23^ (a gift from Eric Kowarz; Addgene # 60515, Cambridge, MA, USA) was digested at SfiI sites (NEB) with the addition of Antarctic Phosphatase (NEB) to prevent re-ligation. Devil *CIITA* was then cloned into SfiI-digested pSBbi-BH using NEBuilder^®^ HiFi DNA Assembly Cloning Kit (NEB). The assembled plasmid pCO2 was transformed into NEB^®^ 5-alpha competent *Escherichia coli* (High Efficiency) (NEB) according to manufacturer’s instructions (see **Supplementary Figure 1** for plasmid maps). Positive clones were identified by colony PCR and the plasmids were isolated using NucleoSpin^®^ Plasmid EasyPure kit (Macherey-Nagel, Düren, Germany). The DNA sequence of the cloned devil *CIITA* transcript was verified by Sanger sequencing using Big Dye^™^ Terminator v3.1 Cycle Sequencing Kit (Applied Biosystems (ABI), Foster City, CA, USA) and Agencourt^®^ CleanSEQ^®^ (Beckman Coulter, Brea, CA, USA) per manufacturer’s instructions. The sequences were analyzed on 3500xL Genetic Analyzer (ABI) (see **Supplementary Table 2** for list of sequencing primers). For detailed step-by-step protocols for plasmid design and construction, reagent recipes, and generation of stable cell lines, see Bio-protocol # e3696^24^.

### 2.3 Transfection and generation of stable cell lines

DFT1 and DFT2 cell line C5065 and JV, respectively, were transfected with plasmid pCO2 to generate stable cell lines that overexpress CIITA. DNA transfections were performed using polyethylenimine (PEI) (1 mg/mL, linear, 25 kDa; Polysciences, Warrington, FL, USA) at a 3:1 ratio of PEI to DNA (w/w) as previously described^8^. Briefly, DFT cells were co-transfected with pCO2 and SB transposase vector pCMV(CAT)T7-SB100^25^ (a gift from Zsuzsanna Izsvak; Addgene plasmid # 34879) at a ratio of 3:1 in μg, respectively. One μg of total plasmid DNA was used per mL of culture volume. The cells were incubated with the transfection solution overnight at 35 °C with 5% CO_2_. The media was removed and replaced with fresh complete RPMI medium. 48 h post-transfection, the cells were observed for expression of reporter gene mTagBFP. Positively-transfected cells were selected with 1 mg/mL hygromycin B (Sigma-Aldrich) for seven days before being maintained in 200 μg/mL hygromycin B in complete RPMI medium. The two tumour cell lines were also transfected with empty vector pSBbi-BH as controls.

### 2.4 RNA sequencing and analysis

RNA libraries were prepared, sequenced and processed as previously described^8,26,27^. **Table 1** shows the source of RNA samples used in this study. Briefly, RNA extraction (two replicates per cell line) was performed using the Nucleospin® RNA Plus Kit (Macherey-Nagel) following the manufacturer’s instructions. mRNA libraries were prepared and sequenced at the Ramaciotti Centre for Genomics (Sydney, NSW, Australia). All RNA samples had RNA Integrity Number (RIN) scores of 10.0. Libraries were prepared using TruSeq Stranded mRNA Library Prep (Illumina Inc., San Diego, CA, USA) and single-end, 100-base pair sequencing were performed on an Illumina NovaSeq 6000 platform (Illumina). The quality of the sequencing reads was assessed using FastQC version 0.11.9^28^. Raw FASTQ files for DFT1.CIITA and DFT2.CIITA have been deposited to the European Nucleotide Archive (ENA) and are available at BioProject # PRJEB45867. Subread version 2.0.0^29^ was used to align sequencing reads to the Tasmanian devil reference genome (GCA_902635505.1 mSarHar1.11) and the number of reads mapped to a gene was counted using featureCounts^30^. The analysis of differentially expressed genes was performed using the statistical software R studio^31^ on R version 4.0.0^32^. Genes with less than 100 aligned reads across all samples were excluded from the analysis and raw library sizes were scaled using *calcNormFactors* in edgeR^33–35^. To account for varying sequencing depths between lanes, read counts were normalised by upper quartile normalisation using *betweenLaneNormalization* in EDASeq^36,37^. Gene length-related biases were normalised by scaling read counts to transcripts per kilobase million (TPM). Differential expression analysis was carried out using the *voom*^38^ function in *limma*^39^ with linear modelling and empirical Bayes moderation^40^. To isolate differentially expressed genes, gene expression of CIITA- or NLRC5-expressing cell lines (DFT.CIITA, DFT.NLRC5) was compared against vector-only control (DFT.BFP) while IFNG-treated cells (DFT.WT + IFNG) was compared against untreated cells (DFT.WT), according to their respective tumour origin. Genes were defined as significantly differentially expressed by applying false discovery rate (FDR) < 0.05, and log_2_ fold change (FC) ≥ 2.0 (upregulated) or ≤ ∼2.0 (downregulated) thresholds (see **Supplementary Table 3** for list of differentially expressed genes).

Venn diagrams of differentially expressed genes were developed using Venny version 2.1^41^. Heatmaps were created from log_2_(TPM) values using the ComplexHeatmap^42^ package in R studio. For functional enrichment analysis, over-representation of gene ontology (GO) biological processes in the list of differentially expressed genes was performed using Database for Annotation, Visualization and Integrated Discovery (DAVID) functional annotation tool^43,44^. The Tasmanian devil *Sarcophilus harrisii* was applied as the species for gene lists and background. Significant GO terms (GOTERM_BP_ALL) were selected by applying the following thresholds: p-value < 0.05 and FDR < 0.05. GO terms were sorted in descending order of fold enrichment values.

To simplify the identification of devil MHC allotypes and maintain consistency in nomenclature to previous works, MHC transcripts in this manuscript were renamed according to Cheng et al., based on sequence similarity^45^ (see **Supplementary Table 4** for corresponding NCBI gene symbols). MHC transcripts *LOC100918485* and *LOC100918744*, which have not been previously characterised, are predicted to encode beta chains of the MHC-II DA gene based on gene homology. These transcripts were renamed as *SAHA-DAB_X1* and *SAHA-DAB_X2*, respectively. Similarly, genes without an official gene symbol (LOC prefixes) were given aliases based on the gene description on NCBI.

### 2.5 Flow cytometric analysis of B2M and MHC-II expression

Cultured cells were harvested using TrypLE™ Express Enzyme (1X) (Thermo Fisher Scientific) and counted using a haemocytometer. 1×10^5^ cells per well were aliquoted into round-bottom 96-well plates and washed with 1X PBS (Thermo Fisher Scientific). Washing steps include centrifugation at 500*g* for 3 min at 4 °C to pellet cells before removal of supernatant. Cells were first stained with Invitrogen™ LIVE/DEAD™ Fixable Near-IR Dead Cell Stain kit (Thermo Fisher Scientific) diluted according to manufacturer’s instructions for 30 min on ice, protected from light. After staining, cells were washed twice with 1X PBS. For MHC-II expression, a monoclonal mouse antibody against the intracellular tail of human HLA-DR α chain was used (Clone TAL.1B5, # M0746, Agilent, Santa Clara, CA, USA). Detection of MHC-I on the surface of cells was performed using a monoclonal mouse antibody against devil beta-2-microglobulin (B2M) in supernatant (Clone 13-34-45; a gift from Hannah Siddle^5^). Cells for intracellular staining of HLA-DR were first fixed and permeabilised using BD Cytofix/Cytoperm™ Plus Fixation/Permeabilization Kit (BD Biosciences, North Ryde, NSW, Australia). All intracellular antibody staining, and washes were carried out in 1X BD Perm/Wash™ Buffer (BD Biosciences) while FACS buffer (PBS with 0.5% BSA, 0.02% sodium azide) was used for surface antibody staining. All cells were incubated with 1% normal goat serum (Thermo Fisher Scientific) for blocking, 10 min on ice. After that, cells were washed and incubated with either anti-human HLA-DRα (0.48 μg/mL) or anti-devil B2M antibody (1:250 v/v dilution) for 30 min on ice. Cells were washed once and stained with goat anti-mouse IgG-Alexa Fluor 488 (2 μg/mL, # A11029, Thermo Fisher Scientific) for 30 min on ice, in the dark. Two final washes were given to remove excess secondary antibody. Fixed cells were resuspended in FACS buffer while the rest were resuspended in FACS fix solution (0.02% sodium azide, 1.0% glucose, 0.4% formaldehyde). Analysis was carried out using Cytek™ Aurora (Cytek Biosciences, Fremont, CA, USA). As a positive control for MHC-I expression, DFT cells were treated with 10 ng/mL devil recombinant IFNG^46^ for 24 h.

### 2.6 Protein extraction and western blot

Cells were harvested and centrifuged at 500*g* for 5 min at room temperature. The pellet was washed twice with cold 1X PBS and weighed. Total cell protein was extracted by adding 1 mL RIPA Lysis and Extraction Buffer (Thermo Fisher Scientific), 10 μL Halt™ Protease Inhibitor Cocktail (Thermo Fisher Scientific) and 10 μL Halt™ Phosphatase Inhibitor Cocktail (Thermo Fisher Scientific) per 40 mg of wet cell pellet. The suspension was sonicated for 30 seconds with 50% pulse and then mixed gently for 15 min on ice. The mixture was centrifuged at 14000*g* for 15 min to pellet the cell debris. The supernatant was transferred to a new tube and total protein was quantified using EZQ® Protein Quantitation kit (Invitrogen) according to manufacturer’s instructions. Two replicates per cell line were prepared for protein extraction. 20 μg of protein per sample was used for target protein detection by western blot. Protein samples were subjected to SDS-PAGE using Bolt™ 4-12%, Bis-Tris, 1.0 mm Mini Protein Gel (Thermo Fisher Scientific). Briefly, protein samples were treated with 1X Bolt™ LDS Sample Buffer (Thermo Fisher Scientific) and 1X Bolt™ Reducing Agent (Thermo Fisher Scientific) at 70 °C for 10 min. Samples were loaded onto the gel and run with 1X Bolt™ MES SDS Running Buffer (Thermo Fisher Scientific) in the Mini Gel Tank (Thermo Fisher Scientific) at 100 V for 5 min followed by 200 V for 15 min. SeeBlue™ Plus2 Pre-stained Protein Standard (Thermo Fisher Scientific) was used as a molecular weight marker. Proteins were transferred to a nitrocellulose membrane using iBlot™ Transfer Stack, nitrocellulose, mini (Thermo Fisher Scientific) and iBlot™ Gel Transfer Device (Thermo Fisher Scientific) using the following settings: 20 V for 7.5 min.

For immunodetection, the membrane was blocked with TBSTM (Tris-buffered saline (TBS): 50 mM Tris-HCl, 150 mM NaCl, pH 7.6), 0.05% Tween 20, and 5% skim milk) for 1 hour at room temperature and rinsed twice with TBST (TBS, 0.05% Tween 20). Then, the membrane was incubated with: (i) rabbit polyclonal anti-beta actin antibody (# ab8227, Abcam, Cambridge, UK) diluted in TBSTM (400 ng/mL), (ii) mouse monoclonal anti-devil SAHA-UA/UB/UC in supernatant (Clone 15-25-18; a gift from Hannah Siddle^10^), or (iii) mouse monoclonal anti-devil SAHA-UK in supernatant (Clone 15-29-1; a gift from Hannah Siddle^10^) overnight at 4 °C. The membranes were washed four times with TBST for a duration of 5 min each wash. After that, the membranes were incubated with either HRP-conjugated goat anti-mouse (250 ng/mL; # P0447, Agilent) or HRP-conjugated goat anti-rabbit immunoglobulin (62.5 ng/mL; # P0448, Agilent) diluted in TBSTM for 1 hour at room temperature. The membranes were given final washes as described above. All incubation and washing steps were performed under agitation. Target protein expression was detected using Immobilon™ Western Chemiluminescent HRP Substrate (Merck Millipore, Burlington, MA, USA) according to manufacturer’s protocol. Protein bands were visualised using Amersham™ Imager 600 (GE Healthcare Life Sciences, Malborough, MA, USA).

### 2.7 Flow cytometric analysis of serum antibody binding

Serum samples from four devils (My, TD4, TD5, and TD6), collected before (pre-immune) and after DFT1 clinical manifestations (immune), were used to assess antibody responses towards CIITA-expressing DFT cell lines (**Supplementary Table 5**). The serum samples were identified as immune from the presence of anti-DFT1 antibodies, which were found to be predominantly against MHC-I on DFT1 cells^8^. ‘My’ was a devil that was immunised, challenged with DFT1 cells, and subsequently treated with an experimental immunotherapy that induced tumour regression^9^. TD4, TD5, and TD6 were naturally DFT1-infected wild devils with either spontaneous tumour regressions (TD4); MHC-II^+^ and CD3^+^ tumour-infiltrating lymphocytes in the tumour (TD5); or B2M^+^ DFT1 cells in fine needle aspirations of a tumour (TD6)^7^. A devil with no clinical signs of DFTD during serum collection (TD7) was included as a negative control for antibody binding towards the DFT cell lines.

Cells were harvested and aliquoted into round-bottom 96-well plates as indicated above. After washing with PBS, cells were stained with LIVE/DEAD™ Fixable Near-IR Dead Cell Stain for 30 min on ice and washed twice with PBS. For blocking, cells were incubated with 1% normal goat serum for 10 min and washed once with FACS buffer. Serum samples were thawed on ice and diluted with FACS buffer (1:50 v/v). 50 μL of serum was added to cells for 1 h and then washed. After that, cells were stained with 10 μg/mL monoclonal mouse anti-devil IgG antibody (A4-D1-2-1, provided by WEHI)^47^ diluted in FACS buffer for 30 min. The cells were washed and stained with 2 μg/mL goat anti-mouse IgG-Alexa Fluor 647 (# A21235, Thermo Fisher Scientific) in FACS buffer for 30 min. After washing, cells were fixed in FACS fix solution and analysed on Cytek™ Aurora. All washing steps include two washes with FACS buffer unless indicated otherwise and all staining steps were carried out on ice, protected from light.

## 3. Results

### 3.1 CIITA plays a dominant role in antigen presentation

To delineate the role of CIITA in DFT cells, differentially expressed genes following stable expression of CIITA were analysed by gene ontology (GO) functional enrichment analysis. Differential expression analysis revealed 888 genes, excluding CIITA, that were modulated (|log_2_FC| ≥ 2, FDR < 0.05) in DFT1.CIITA compared to vector-only cell line DFT1.BFP (**Figure 1**, **Supplementary Table 3**). In DFT2.CIITA, there were 56 genes that were differentially expressed relative to DFT2.BFP. Ten genes were commonly up- or down-regulated by CIITA in DFT1 and DFT2 cells. Most of these genes were of the MHC-II antigen processing and presentation pathway. *SAHA-DAA, SAHA-DAB2* and *SAHA-DAB3* are devil classical MHC-II genes while *SAHA-DMA* and *SAHA-DMB* encode non-classical MHC-II. Others include *CD74* and gamma-interferon-inducible lysosomal thiol reductase (*IFI30*), which encode the invariant chain, an MHC-II chaperone, and an enzyme for lysosomal degradation of proteins, respectively. Except for *IFI30*, these genes were among the most highly upregulated genes in the transcriptome of DFT1.CIITA (**Table 2**) and DFT2.CIITA (**Table 3**).

**Figure 1.**
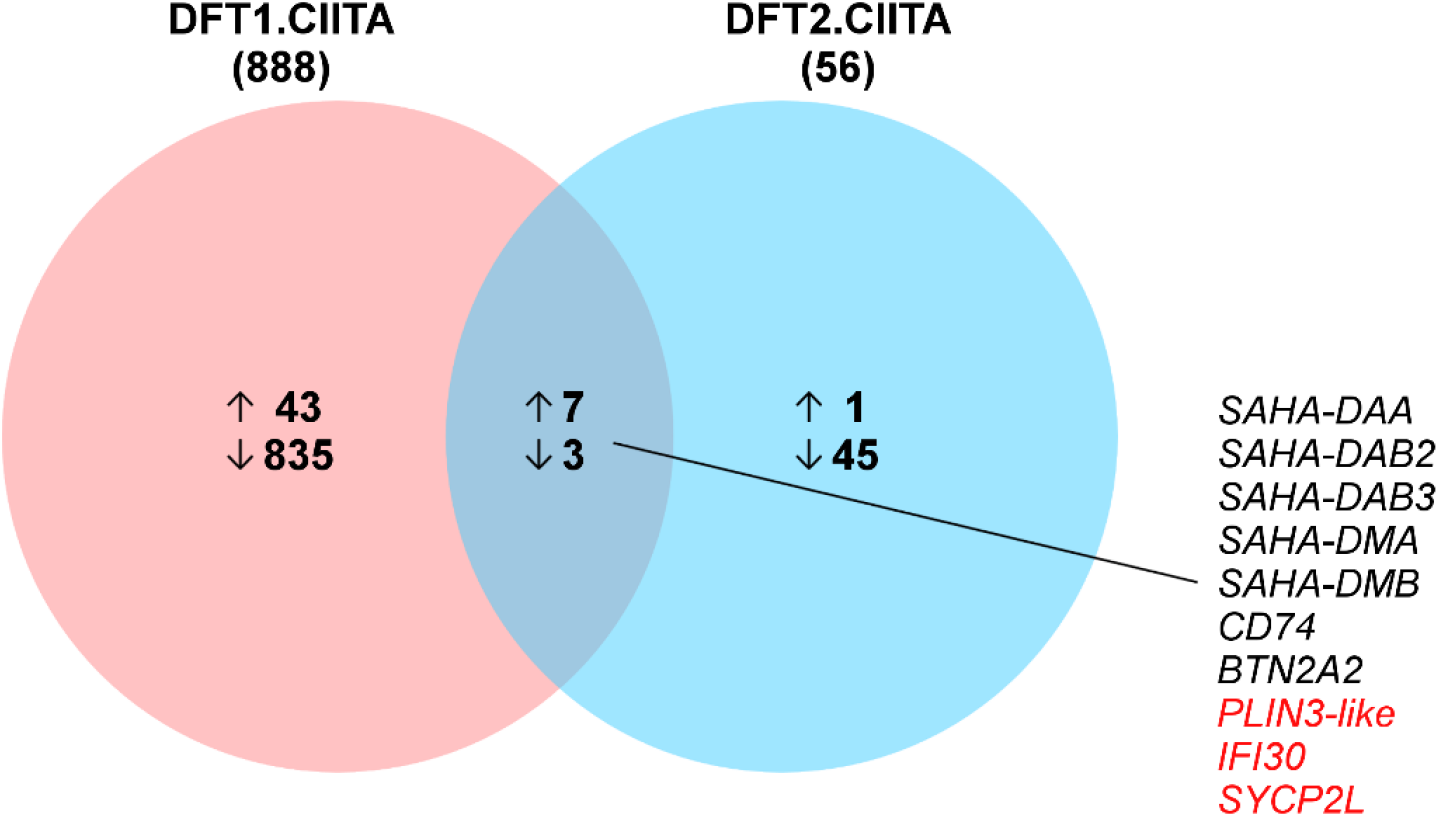
Venn diagram of differentially expressed genes in DFT1 and DFT2 cells with CIITA overexpression. Change in gene expression was identified between DFT.CIITA and vector-only control DFT.BFP. Total number of DEGs is indicated in parenthesis under sample name and the number of upregulated (↑) and downregulated genes (↓) are described in each Venn circle. Mutually inclusive genes that were upregulated are indicated in black and downregulated in red.

**Table 2.** Top 20 most significantly upregulated genes in DFT1.CIITA

| Gene | Gene description | MHC pathway | log<sub>2</sub>FC | FDR |
| --- | --- | --- | --- | --- |
| *SAHA-DAA* | Class II histocompatibility antigen, DA alpha chain | Class II | 17.09 | 1.90E-04 |
| *CD74* | CD74 molecule | Class II | 16.39 | 7.77E-05 |
| *CIITA* | Class II major histocompatibility complex transactivator | Class II | 15.40 | 1.07E-04 |
| *SAHA-DMB* | Class II histocompatibility antigen, DM beta chain | Class II | 10.26 | 7.00E-05 |
| *SAHA-DAB_X2* | Class II histocompatibility antigen, DA beta chain | Class II | 9.04 | 1.11E-03 |
| *SAHA-DAB_X1* | Class II histocompatibility antigen, DA beta chain | Class II | 8.82 | 7.77E-04 |
| *PSMB9* | Proteasome 20S subunit beta 9 | Class I | 8.52 | 4.08E-04 |
| *SAHA-DMA* | Class II histocompatibility antigen, DM alpha chain | Class II | 6.52 | 5.12E-04 |
| *TAP1* | Transporter 1, ATP binding cassette subfamily B member | Class I | 6.46 | 6.95E-04 |
| *PSMB8* | Proteasome 20S subunit beta 8 | Class I | 6.13 | 1.29E-03 |
| *SAHA-DAB3* | Class II histocompatibility antigen, DA beta chain | Class II | 6.08 | 7.72E-05 |
| *SAHA-UC* | Class I histocompatibility antigen heavy chain | Class I | 5.08 | 2.17E-03 |
| *SAHA-UA* | Class I histocompatibility antigen heavy chain | Class I | 4.77 | 2.29E-03 |
| *B2M* | Beta-2-microglobulin | Class I | 4.43 | 1.96E-05 |
| *SAHA-UB* | Class I histocompatibility antigen heavy chain | Class I | 4.41 | 2.25E-03 |
| *SAHA-DAB2* | Class II histocompatibility antigen, DA beta chain | Class II | 4.21 | 1.96E-05 |
| *ICOSLG* | Inducible T Cell Costimulator (ICOS) Ligand | Unrelated | 3.98 | 1.14E-03 |
| *KIF6* | Kinesin family member 6 | Unrelated | 3.88 | 1.10E-02 |
| *BARX1* | BARX homeobox 1 | Unrelated | 3.68 | 4.99E-03 |
| *MID1* | Midline 1 | Unrelated | 3.67 | 3.52E-03 |

See **Supplementary Table 3** full list of differentially expressed genes and log<sub>2</sub>TPM values.

**Table 3.** Significantly upregulated genes in DFT2.CIITA

| Gene | Gene description | MHC pathway | log<sub>2</sub>FC | FDR |
| --- | --- | --- | --- | --- |
| *CD74* | CD74 molecule | Class II | 13.60 | 6.69E-03 |
| *CIITA* | Class II major histocompatibility complex transactivator | Class II | 11.50 | 2.31E-03 |
| *SAHA-DAA* | Class II histocompatibility antigen, DA alpha chain | Class II | 8.52 | 2.31E-03 |
| *SAHA-DMB* | Class II histocompatibility antigen, DM beta chain | Class II | 8.34 | 2.81E-04 |
| *SAHA-DAB3* | Class II histocompatibility antigen, DA beta chain | Class II | 7.09 | 2.19E-03 |
| *SAHA-DMA* | Class II histocompatibility antigen, DM alpha chain | Class II | 6.82 | 6.82E-04 |
| *SAHA-DAB2* | Class II histocompatibility antigen, DA beta chain | Class II | 4.14 | 3.10E-04 |
| *BTN2A2* | Butyrophilin subfamily 2 member A2 | Unrelated | 3.49 | 3.54E-02 |
| *NDUFA4L2* | NDUFA4 mitochondrial complex associated like 2 | Unrelated | 2.83 | 2.72E-02 |

See **Supplementary Table 3** full list of differentially expressed genes and log<sub>2</sub>TPM values.

In DFT1.CIITA, several MHC-I heavy chain and accessory genes were strongly induced, depicting a role of CIITA in MHC-I antigen presentation (**Table 2**). These include: (i) MHC-I heavy alpha chain genes *SAHA-UA, SAHA-UB*, and *SAHA-UC*; and (ii) *B2M*, which associates with MHC-I alpha chains to form the trimeric structure of MHC-I molecules; (iii) transporter associated with antigen processing 1 (*TAP1*) for peptide transport into the endoplasmic reticulum; and (iv) proteasomal subunits *PSMB8* and *PSMB9*.

Next, all significantly up- or down-regulated genes were analysed for enriched GO biological processes using DAVID bioinformatics resource. Thresholds *p* value < 0.05 and FDR < 0.05 were applied to filter out insignificant over-represented GO terms. The most significantly enriched GO biological process in the list of upregulated genes in DFT1.CIITA and DFT2.CIITA was *antigen processing and presentation* (GO:0019882) followed by *immune response* (GO:0006955) (**Table 4**). Both processes were identified in genes of the MHC-I and MHC-II machinery (**Supplementary Tables 6** and **7**). C*ell adhesion* (GO:0007155) and *cell communication* (GO:0007154) were enriched in genes downregulated in DFT1.CIITA; there were no GO biological processes that were associated with downregulated genes in DFT2.CIITA.

**Table 4.** GO biological processes enriched in differentially expressed genes in DFT1.CIITA and DFT2.CIITA

| GO ID | GO term | Count | Term size | Fold enrichment | *p* value | FDR |
| --- | --- | --- | --- | --- | --- | --- |
| **DFT1.CIITA** |  |  |  |  |  |  |
| *Upregulated* |  |  |  |  |  |  |
| GO:0019882 | antigen processing and presentation | 8 | 42 | 86.09 | 1.34E-12 | 1.19E-09 |
| GO:0006955 | immune response | 9 | 518 | 7.85 | 4.75E-06 | 2.12E-03 |
| *Downregulated* |  |  |  |  |  |  |
| GO:0007155 | cell adhesion | 44 | 719 | 2.14 | 2.27E-06 | 4.48E-03 |
| GO:0022610 | biological adhesion | 44 | 721 | 2.13 | 2.44E-06 | 4.48E-03 |
| GO:0023052 | signaling | 113 | 2746 | 1.44 | 4.94E-06 | 6.05E-03 |
| GO:0044700 | single organism signaling | 111 | 2726 | 1.42 | 1.12E-05 | 1.02E-02 |
| GO:0007154 | cell communication | 112 | 2773 | 1.41 | 1.46E-05 | 1.07E-02 |
| **DFT2.CIITA** |  |  |  |  |  |  |
| *Upregulated* |  |  |  |  |  |  |
| GO:0019882 | antigen processing and presentation | 4 | 42 | 215.21 | 9.33E-08 | 3.90E-05 |
| GO:0006955 | immune response | 4 | 518 | 17.45 | 1.87E-04 | 3.91E-02 |

### 3.2 Regulation of MHC-I and MHC-II pathway by CIITA

To further characterise the regulation of MHC-I and MHC-II by CIITA and how it differs from IFNG or NLRC5 stimulation, a heatmap was used to display the relative expression of MHC-I and MHC-II genes, and key accessory proteins between the different treatments. The transcriptome of IFNG-treated DFT2 cells was previously carried out on DFT2 cell line RV (DFT2.WT^RV^)^27^ while subsequent experiments on DFT2 cells were performed using DFT2 cell line JV (DFT2.WT). Schwann cell differentiation marker SRY-box 10 (*SOX10*) and neuroepithelial marker nestin (*NES*) were used as internal gene controls, and myelin protein periaxin (*PRX*) was used to discriminate DFT1 cells from DFT2.

As described above, CIITA induced the transcription of *B2M;* MHC-I heavy chains *SAHA-UA, -UB, -UC*; *PSBM8; PSMB9;* and *TAP1* in DFT1 cells. There was also an upregulation of non-classical MHC-I *SAHA-UK*, and downregulation of *NLRC5* and proteasomal subunit *PSBM10* in DFT1.CIITA cells (**Figure 2**). Excluding *NLRC5*, genes that were modulated in DFT1.CIITA were synonymously up- or down-regulated in DFT1.NLRC5, suggesting similar roles of CIITA to NLRC5 in DFT1 cells. However, induction of the MHC-I pathway by CIITA was not as strong as NLRC5 despite having similarly high levels of expression in the respective cell lines (**Figure 2**, **Supplementary Table 3**). IFNG exhibited a wider range in regulation of genes from the MHC-I pathway compared to NLRC5 and CIITA. Peptide transporter *TAP2* and MHC-I chaperone TAP binding protein (*TAPBP*) were exclusively upregulated by IFNG in DFT1 and DFT2 cells. Meanwhile, the expression of CIITA in DFT2 cells did not appear to significantly influence any of the MHC-I machinery.

**Figure 2.**
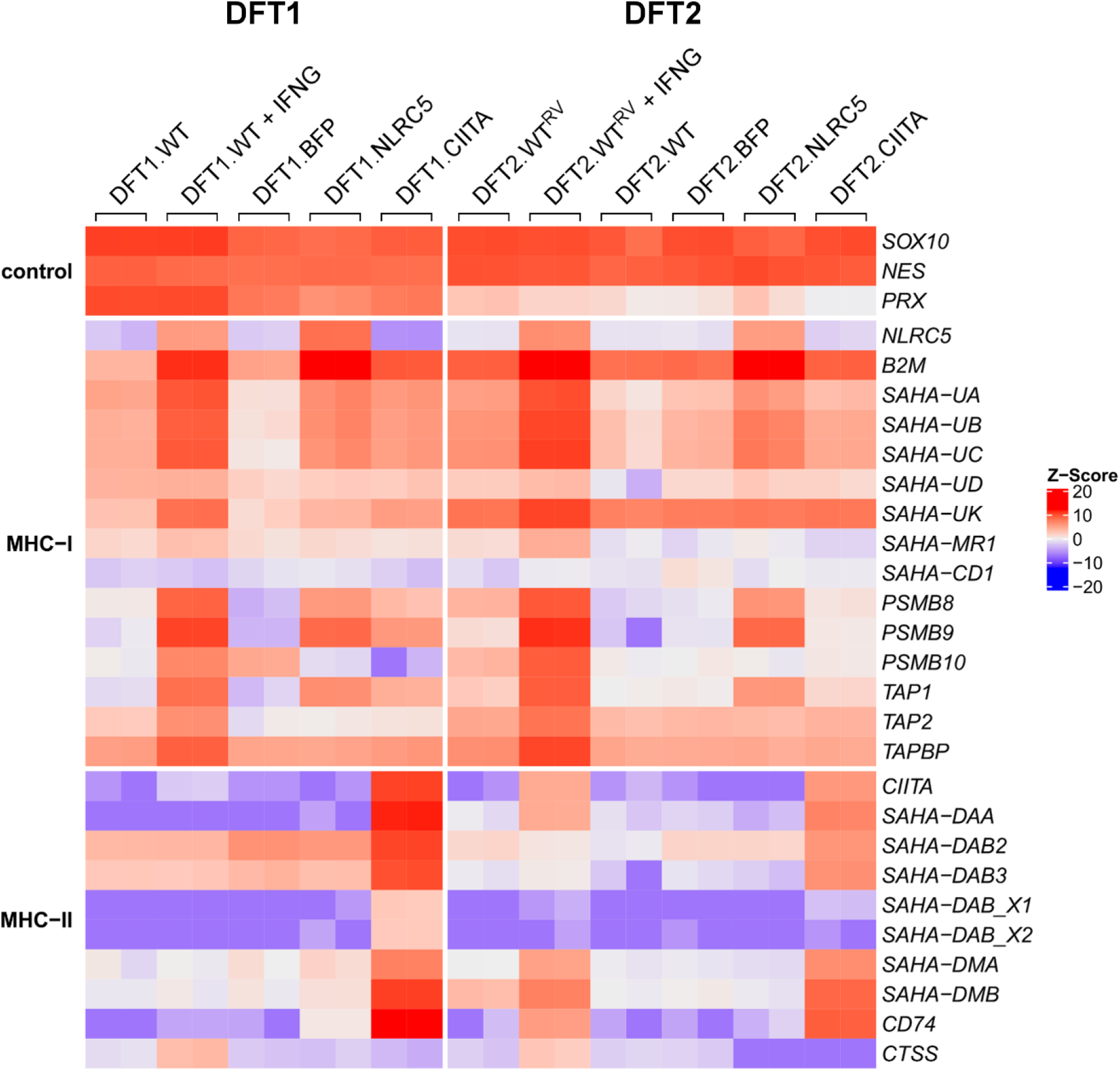
Heatmap showing relative expression of genes involved in MHC-I and MHC-II antigen processing and presentation in wild type, IFNG-treated, BFP- (vector control), NLRC5-, and CIITA-expressing DFT1 and DFT2 cells. Z-scores were calculated from log_2_TPM expression values and scaled across each gene (rows). High and low relative expression are represented by red and blue, respectively. Replicates per treatment (*N*=2) are included in the heatmap.

High levels of CIITA transcripts in DFT1.CIITA was correlated with strong induction of all the MHC-II genes, with *SAHA-DAB_X1* and *SAHA-DAB_X2* being the weakest. This was not observed in the other cell lines nor in DFT2.CIITA. CIITA was expressed to a lesser extent in DFT2.CIITA relative to DFT1.CIITA, and all MHC-II genes but *SAHA-DAB_X1* and *SAHA-DAB_X2* were upregulated. The expression of *CIITA*, MHC-II genes and *CD74* was relatively low in DFT1.WT and DFT2.WT cells except for *SAHA-DAB2* and *SAHA-DAB3* in DFT1.WT. There was a moderate increase in CIITA expression after IFNG treatment in DFT1 cells, but it was insufficient to initiate transcription of MHC-II genes or *CD74*. In IFNG-treated DFT2 cells where CIITA was induced to a higher degree, there was only partial activation of the MHC-II gene set (*SAHA-DAA, SAHA-DMA, SAHA-DMB*), and an upregulation of *CD74*. Interestingly, MHC-II protease cathepsin *CTSS* was only induced with IFNG treatment in DFT1 and DFT2 cells.

### 3.3 MHC-I and MHC-II molecules are upregulated by CIITA in DFT cells

MHC-II (HLA-DRA) protein expression was absent in wild type (WT) DFT1 and DFT2 cells and in vector-only transfected cells (BFP) but was significantly upregulated in CIITA-expressing DFT1 cells (**Figure 3A**). In DFT2 cells, the overexpression of CIITA did not alter median MHC-II expression, or more specifically MHC-II gene loci HLA-DRA. Neither IFNG treatment nor NLRC5 overexpression induced MHC-II protein expression in DFT1 and DFT2 cells. CIITA was capable of restoring surface expression of B2M in DFT1 cells, albeit to a lesser degree than NLRC5 and IFNG stimulation, consistent with the transcriptomic results (**Figure 3A, Figure 2**). Meanwhile the basal expression of B2M in DFT2 cells was enhanced slightly by CIITA.

**Figure 3.**
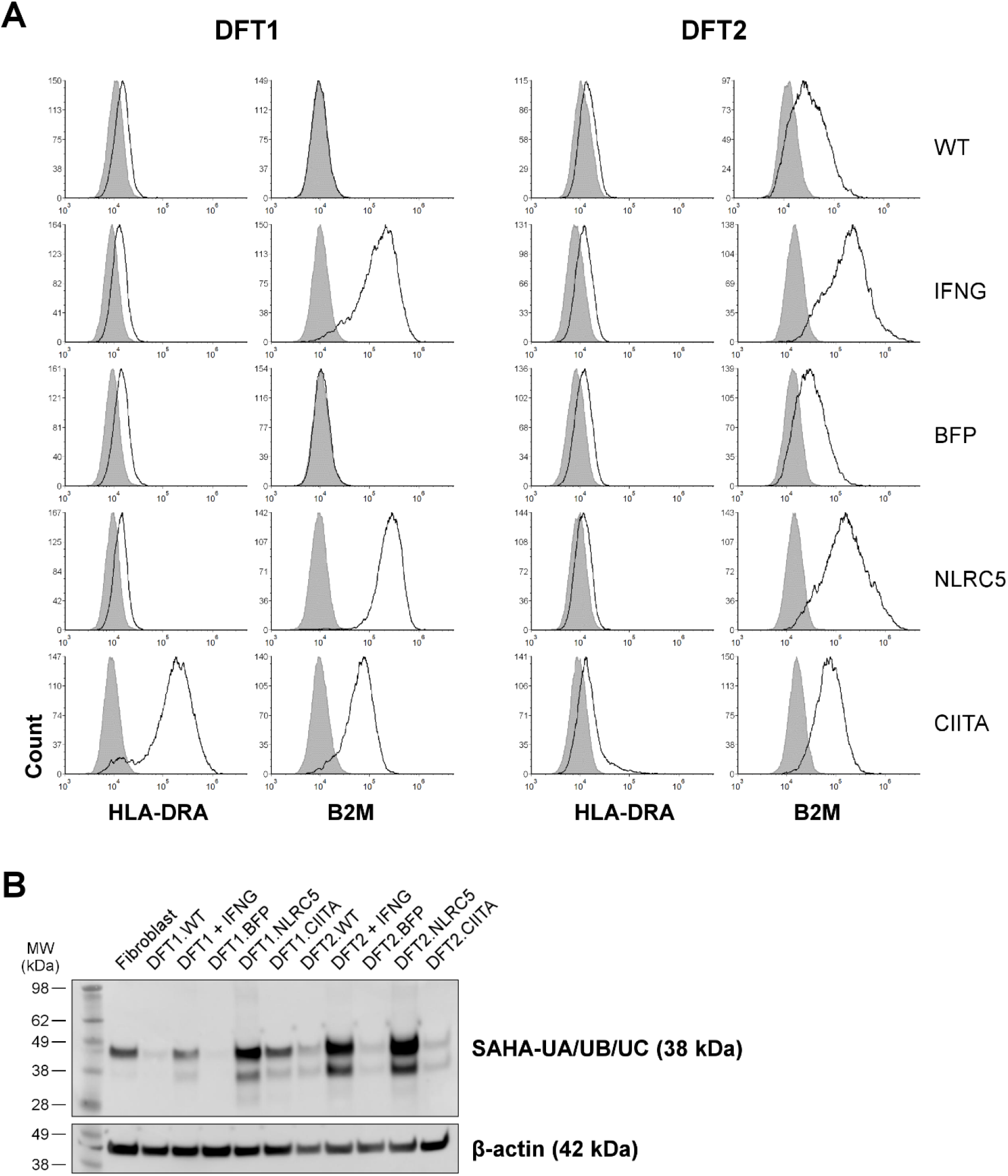
Expression of MHC-II, B2M and MHC-I in DFT1 and DFT2 cell lines. (**A**) Wild type (WT), IFNG-treated (IFNG), vector-only control (BFP), NLRC5-overexpressing (NLRC5) or CIITA-overexpressing (CIITA) DFT1 and DFT2 cells were analysed by flow cytometry for B2M and MHC-II expression using antibodies against surface devil B2M or intracellular HLA-DR alpha chain (HLA-DRA), respectively (*solid line*). B2M and MHC-II expressions were overlaid with a secondary antibody-only control (*shaded area*). The results shown are representative of *N*=3 replicates/treatment. (**B**) Cell lysate from devil fibroblast, DFT1 and DFT2 cell lines was incubated with an antibody against MHC-I heavy chain genes SAHA-UA/UB/UC for western blot analysis of MHC-I expression. β-actin was included as a loading control. *MW*, molecular weight.

In agreement with an increase in surface B2M expression on DFT1.CIITA by flow cytometry, an upregulation of MHC-I heavy chains was detected by western blot compared to wild type (DFT1.WT) and vector-only cells (DFT1.BFP) (**Figure 3B**). IFNG-treated and NLRC5-overexpressing DFT1 and DFT2 cells also expressed elevated levels of MHC-I heavy chains. Although flow cytometry detected an increase in B2M expression on DFT2.CIITA, the expression of MHC-I heavy chains by western blot was similar to DFT2.WT and DFT2.BFP.

### 3.4 Analysis of anti-DFT serum antibody response against CIITA-induced antigens

We have previously shown that MHC-I on DFT1 cells is the predominant antibody target in devils with natural and induced anti-DFT immune response including tumour regressions^8^. Here we tested if expression of CIITA in DFT cells could also upregulate antibody targets on DFT cells. Four devils (My, TD4, TD5, TD6) that developed DFT1 tumours and subsequent serum antibodies (immune) that bound MHC-I were selected for screening against CIITA-expressing DFT1 and DFT2 cells. Serum from each devil prior to DFT1 infection or observable DFT1 tumours (pre-immune) was included to assess the change in antibody levels after DFT1 infection.

Relative to MHC-I negative DFT1.WT and DFT1.BFP, serum antibodies from all four devils post-DFT1 development generally showed higher binding to DFT1 cells overexpressing NLRC5 (**Figure 4**). Antibody levels against CIITA-expressing DFT1 cells were higher than DFT1.WT and DFT1.BFP in immune sera from My, a captive devil with an immunotherapy-induced DFT1 regression, and TD4, a wild devil with a natural DFT1 regression. Binding of serum antibodies to DFT1.CIITA cells was relatively lower than DFT1.NLRC5. There was no increase in antibody binding towards DFT1.CIITA compared to DFT1.WT and DFT1.BFP from immune sera of devils TD5 and TD6.

**Figure 4.**
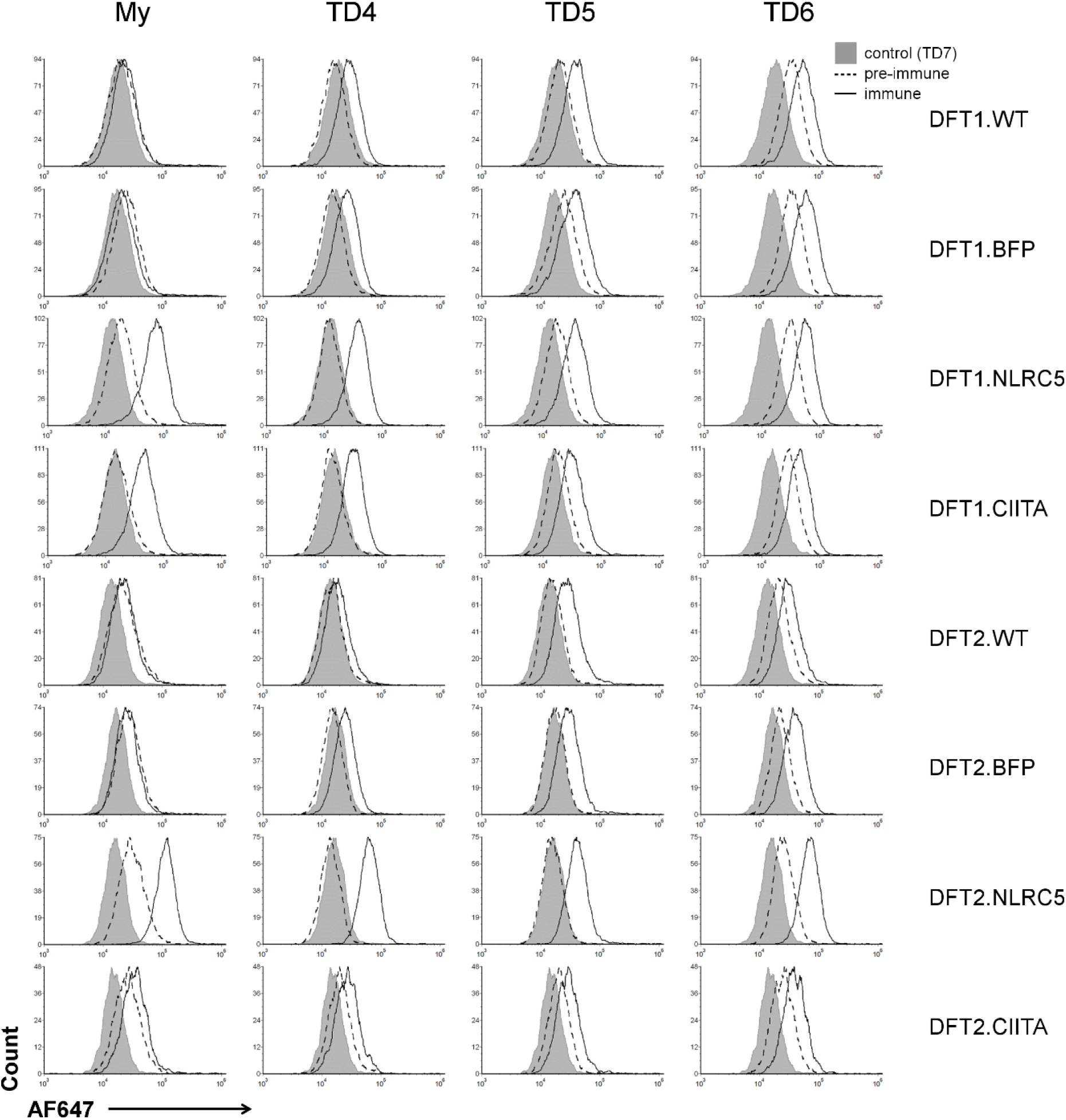
Flow cytometric analysis of serum antibody response towards DFT1 and DFT2 cells overexpressing CIITA. Sera from four devils (My, TD4, TD5, TD6) with antibody responses to MHC-I^+^ DFT1 cells after DFT1 infection (immune) were used. Antibody binding was compared against wild type (DFT.WT), vector-only (DFT.BFP) and NLRC5-overexpressing cells (DFT.NLRC5). Serum collected prior to infection (pre-immune) and from a non-infected devil (TD7) were included as negative controls. *AF647*, Alexa Fluor 647.

Serum from DFT1-infected devils reacted with DFT2 cells but only following NLRC5 overexpression. Serum from My, TD4, TD5 and TD6 all had strong antibody binding to DFT2.NLRC5 which was not observed in the other DFT2 cell lines. This suggests that NLRC5 upregulates similar antigenic target(s) in DFT1 and DFT2 cells.

## 4. Discussion

Clonally transmissible cancers in nature are rare, and yet the Tasmanian devils are affected by two of the only three known naturally occurring transmissible cancers in vertebrates. In a cancer where allogeneity exists between individual host tissues and tumour, allogeneic MHC molecules on tumour cells are important targets for anti-tumour immunity. MHC-I expression on DFT1 cells has been exploited for vaccine development and immunotherapy to enhance anti-DFT immunity via CD8^+^ T cell responses^9^. In this study, we showed that the class II transactivator CIITA can modulate MHC-I and MHC-II antigen processing and presentation pathways in DFT cells. Surprisingly, the overexpression of CIITA resulted in upregulation of MHC-I and MHC-II molecules in DFT1 cells but not DFT2 cells.

MHC-II expression is normally confined to a subset of haematopoietic antigen-presenting cells, and DFT1 and DFT2 cells do not typically express MHC-II genes and proteins. We demonstrated the expression of MHC-II proteins in non-haematopoietic DFT1 cells through CIITA-induced upregulation of classical and non-classical MHC-II genes, and the invariant chain *CD74*. The lack of detectable MHC-II proteins in CIITA-expressing DFT2 cells could be due to insufficient expression of MHC-II genes and *CD74* for stable expression of MHC-II molecules. Post-transcriptional regulation might be involved in MHC-II expression in DFT cells, as described in human T cells^48^. However, the regulation of MHC-II expression by CIITA in a quantitative (and qualitative) manner, in which CIITA is the rate-determining factor for mRNA and protein expression of MHC-II genes^49^, suggests a correlation between lack of MHC-II protein expression and the relatively low CIITA expression in DFT2.CIITA compared to DFT1.CIITA. A heterozygous non-synonymous mutation (D59N) in transcription factor *RFX5* is present in DFT2 tumours^6^. RFX5 is a transcription factor of the multiprotein MHC enhanceosome that regulates MHC-I and MHC-II expression^50,51^. Although transcription of MHC-I and MHC-II genes were inducible in DFT2 cells following stimulation, the functional impact of this mutation on MHC transcription remains to be explored.

Differential expression of MHC-II allotypes upon CIITA induction, as observed with *SAHA-DAB_X1* and *SAHA-DAB_X2* that were consistently expressed at lower levels compared to other MHC-II genes, suggests additional regulatory mechanism(s) that control the expression of MHC-II genes beyond that of CIITA. Variations in expression levels of MHC-I and MHC-II genes have been associated with sequence polymorphism in the promoter or 3’ untranslated region (UTR) of MHC genes, which modulates transcription either epigenetically or non-epigenetically, in addition to post-transcriptional regulation^52–54^. The varying degrees of inducibility and expression of devil MHC-II allotypes could correlate to tissue-specific expression, with functions that differ from classical MHC-II genes.

Consistent with findings from pioneering studies on CIITA function^20,21^, CIITA exhibited transcriptional activity over the MHC-I pathway in DFT1 cells that lack MHC-I expression. The ability of CIITA to regulate MHC-I expression is attributed to similarities in the regulatory elements at the proximal promoters of MHC-I and MHC-II genes, and interaction with the same transcription factors of the MHC enhanceosome as NLRC5^20,21,51,55,56^. In MHC-I positive DFT2 cells, overexpression of CIITA resulted in minimal upregulation of MHC-I compared with NLRC5 or IFNG stimulation. The limited CIITA influence on MHC-I expression is commonly observed in cells with high constitutive levels of MHC-I^20,21^. This illustrates the role of NLRC5 as the primary transactivator for MHC-I expression and a secondary role for CIITA.

Unlike the ubiquitous expression of MHC-I molecules in nucleated cells, MHC-II expression is tightly regulated in a cell type-, differentiation-, and stimulus-specific manner. Evidence for inducibility of MHC-II expression in DFT cells suggests that MHC-II-restricted tumour antigen presentation could occur in the physiological setting under inflammatory conditions that upregulate CIITA. This could provide additional targets for allogeneic antibody responses, as our results show that CIITA upregulation increases binding of serum antibodies collected from devils that had both natural and immunotherapy-induced DFT1 regressions. In canine transmissible venereal tumour (CTVT), the tumour regression phase is often associated with upregulation of MHC-I and MHC-II molecules, mediated by factors such as IFNG from tumour infiltrating lymphocytes^57,58^.

The capacity to express MHC-II molecules with CIITA expression could stem from the Schwann cell origins of DFT1 and DFT2 cells^27,59^. Schwann cells express MHC-II molecules upon traumatic and inflammatory injury, playing a role in antigen presentation to CD4^+^ T cells to modulate local immune responses^60,61^. Similarly, CIITA-expressing DFT cells have the potential to present MHC-II-restricted tumour antigens to CD4^+^ T cells and potentiate anti-DFT immune responses. Several studies in murine models have demonstrated immune-mediated tumour rejection and/or tumour growth retardation using MHC-II-expressing tumour cell lines, either through CIITA or MHC-II gene transfer^62–67^. These primary responses were also protective against subsequent challenge with parental MHC-II negative tumours. The expression of MHC-II on CIITA-expressing DFT cells can offer insight into the importance of CD4^+^ T cells in the interplay with other immune cells for anti-tumour immunity and allograft rejection.

In this study, the role of CIITA as the master regulator of MHC-II expression was reaffirmed in a non-model immunology research species. We have delineated the regulation of MHC-I and MHC-II pathways by CIITA in marsupials and transmissible cancers. The ability to induce MHC-II expression in transmissible tumour cells creates an avenue for vaccine and immunotherapeutic strategies to enhance anti-tumour immunity through CD4^+^ T cell help and inform of the importance of MHC-II in anti-tumour and allogeneic immune responses. The relatively simple process we developed for making cell lines that constitutively express NLRC5 and CIITA can be readily adapted for many other species and potentially be used in conjunction with CD80/CD86 to provide antigen stimulation in *in vitro* assays. This is critical for 99% of species that lack reagents, such as agonistic anti-CD3 and anti-CD28 antibodies, for inducing T cell activation *in vitro*.

## Authors’ contributions

ABL, ASF, CEBO and GMW designed the study. ASF and CEBO developed the technology. CEBO performed the experiments and bioinformatic analyses. CEBO created the figures. ABL, ASF, CEBO, GMW, HVS and YC analysed and interpreted the data. CEBO wrote the manuscript, and all authors edited the manuscript.

## Acknowledgements

The authors would like to thank Jocelyn Darby and Alana De Luca for assistance in the lab, Terry Pinfold for assistance in flow cytometry, and Amanda Patchett for assistance in bioinformatic analysis. We wish to thank G. Ralph for ongoing care of Tasmanian devils, the Bonorong Wildlife Sanctuary for providing access to Tasmanian devils, and R. Pye for providing care for devils and collecting blood samples.

