## Supplementary Materials for "CIITA induces expression of MHC-I and MHC-II in transmissible cancers"

**Andrew S. Flies, PhD**

Menzies Institute for Medical Research, College of Health and Medicine
University of Tasmania
Private Bag 23, Hobart TAS 7000

Supplementary Figure 1. Schematic diagram of plasmids pSBbi-BH, pCO1 and pCO2

Supplementary Table 1. PCR primers and reaction conditions

Supplementary Table 2. Sequencing primers and reaction conditions

Supplementary Table 3. Differential expression analysis [*separate file*]

Supplementary Table 4. List of gene aliases used in this study and their corresponding NCBI gene symbol

Supplementary Table 5. Serum sample information

Supplementary Table 6. Gene ontology (GO) biological processes enriched in DFT1.CIITA [*separate file*]

Supplementary Table 7. Gene ontology (GO) biological processes enriched in DFT2.CIITA [*separate file*]

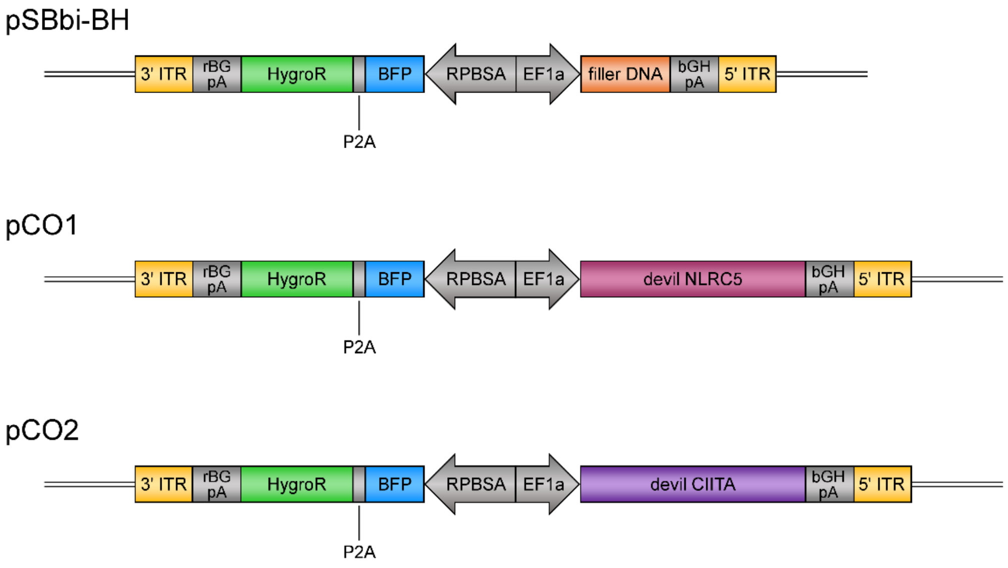


**Supplementary Figure 1**. Schematic diagram of plasmids pSBbi-BH, pCO1 and pCO2. *ITR*, inverted terminal repeats; *pA*, polyadenylation tail; *HygroR*, hygromycin resistance gene; *BFP*, mTagBFP.

**Supplementary Table 1**. PCR primers and reaction conditions

| Vector | Template DNA | Target DNA | Product size (bp) | Primers | Primer sequence (5' to 3') | Reaction conditions |
|---|---|---|---|---|---|---|
| pCO2 | pSBbi-BH | SfiI-digested pSBbi-BH | 6839 | N/A | N/A | N/A |
| | devil PBMC cDNA | devil *CIITA* cDNA (XM_023497584.2) | 3360 | pCO2_X2.FOR | GAAAACTACCCCAAGCTGGCCTCTGAGGCCGCCACCATGCGGCAGTTACGTGGTGC | 1x: 98 °C for 30 s;<br>10x: 98 °C for 10 s, 65 to 56 °C (-1 °C each cycle) for 30 s, 72 °C for 3 min;<br>25x: 98 °C for 10 s, 68 °C for 30 s, 72 °C for 3 min;<br>1x: 72 °C for 2 min |
| | | | | pCO2.REV | GAGAATTGATCCCCAAGCTTGGCCTGACAGGCCTTATCGCAAGCTTATTCGGGAGTCC | |

**Supplementary Table 2**. Sequencing primers and reaction conditions

| Template DNA | Target DNA | Primers | Primer sequence (5' to 3') | Reaction conditions |
|---|---|---|---|---|
| pCO2 | devil *CIITA* cDNA | pSB_EF1a_seq.FOR | GCCTCAGACAGTGGTTCAAAG | 1x: 96°C for 1 min;<br>25x: 96°C for 10 s, 50°C for 5 s, 60°C for 4 min |
| | | deCIITA_seq1.FOR | GACAGTTGGGAGACCAAACG | |
| | | deCIITA_seq2.FOR | AGTACTGGGCAAGGCTGGTC | |
| | | deCIITA_seq3.FOR | CATCTGAAACTTGGTCCAGAGAC | |
| | | deCIITA_seq4.FOR | TGCTCAGTCACTGCTACAGC | |
| | | CIITAF | ACCCTTGTCCAACTTGGTTGTGTTACC | |
| | | pSB_bGH_seq.REV | AGGCACAGTCGAGGCTGAT | |

**Supplementary Table 4**. List of gene aliases used in this study and their corresponding NCBI gene symbol

| Gene | NCBI gene symbol | Gene description | MHC pathway |
|---|---|---|---|
| *SAHA-UA* | *SAHAI-01* | Class I histocompatibility antigen heavy chain | Class I |
| *SAHA-UB* | *LOC100927947* | Class I histocompatibility antigen heavy chain | Class I |
| *SAHA-UC* | *LOC105750614* | Class I histocompatibility antigen heavy chain | Class I |
| *SAHA-UD* | *SAHAI-12* | Class I histocompatibility antigen heavy chain | Class I |
| *SAHA-UK* | *LOC100926320* | Class I histocompatibility antigen heavy chain | Class I |
| *SAHA-MR1* | *LOC100917175* | Class I histocompatibility antigen heavy chain | Class I |
| *SAHA-CD1* | *LOC100928203* | Class I histocompatibility antigen heavy chain | Class I |
| *SAHA-DAA* | *LOC100923003* | Class II histocompatibility antigen, DA alpha chain | Class II |
| *SAHA-DAB2* | *LOC100930980* | Class II histocompatibility antigen, DA beta chain | Class II |
| *SAHA-DAB3* | *LOC100922474* | Class II histocompatibility antigen, DA beta chain | Class II |
| *SAHA-DAB_X1* | *LOC100918485* | Class II histocompatibility antigen, DA beta chain | Class II |
| *SAHA-DAB_X2* | *LOC100918744* | Class II histocompatibility antigen, DA beta chain | Class II |
| *SAHA-DMA* | *LOC100925801* | Class II histocompatibility antigen, DM alpha chain | Class II |
| *SAHA-DMB* | *LOC100925533* | Class II histocompatibility antigen, DM beta chain | Class II |
| *BTN2A2* | *LOC100930971* | Butyrophilin subfamily 2 member A2 | Unrelated |
| *PLIN3-like* | *LOC116419077* | Perilipin-3-like | Unrelated |
| *ICOSLG* | *LOC100929908* | Inducible T Cell Costimulator (ICOS) Ligand | Unrelated |

**Supplementary Table 5**. Serum sample information

| Devil name | Devil ID | Sample collection date | |
|---|---|---|---|
| | | **Pre-immune** | **Immune** |
| My | Missy | 10 Mar 2011 | 3 Jan 2013 |
| TD4 | WPP314 | July 2011 | May 2014 |
| TD5 | WPP209 | Nov 2010 | May 2011 |
| TD6 | WPP240 | Nov 2013 | Feb 2014 |
| TD7 | WPP213 | Feb 2012 | N/A |

*TD4–TD6 correspond to devils used in* Pye et al. (2016) *and* Ong et al. (2021)
